# Wx: a neural network-based feature selection algorithm for next-generation sequencing data

**DOI:** 10.1101/221911

**Authors:** Sungsoo Park, Bonggun Shin, Yoonjung Choi, Kilsoo Kang, Keunsoo Kang

## Abstract

**Motivation:** Next-generation sequencing (NGS), which allows the simultaneous sequencing of billions of DNA fragments simultaneously, has revolutionized how we study genomics and molecular biology by generating genome-wide molecular maps of molecules of interest. For example, an NGS-based transcriptomic assay called RNA-seq can be used to estimate the abundance of approximately 190,000 transcripts together. As the cost of next-generation sequencing sharply declines, researchers in many fields have been conducting research using NGS. The amount of information produced by NGS has made it difficult for researchers to choose the optimal set of target genes (or genomic loci).

**Results:** We have sought to resolve this issue by developing a neural network-based feature (gene) selection algorithm called Wx. The Wx algorithm ranks genes based on the discriminative index (*DI*) score that represents the classification power for distinguishing given groups. With a gene list ranked by *DI* score, researchers can institutively select the optimal set of genes from the highest-ranking ones. We applied the Wx algorithm to a TCGA pan-cancer gene-expression cohort to identify an optimal set of gene-expression biomarker (universal gene-expression biomarkers) candidates that can distinguish cancer samples from normal samples for 12 different types of cancer. The 14 gene-expression biomarker candidates identified by Wx were comparable to or outperformed previously reported universal gene expression biomarkers, highlighting the usefulness of the Wx algorithm for next-generation sequencing data. Thus, we anticipate that the Wx algorithm can complement current state-of-the-art analytical applications for the identification of biomarker candidates as an alternative method.

**Availability:** https://github.com/deargen/DearWX

**Supplementary information:** Supplementary data are available at online.

## 1 Introduction

Advances in science and technology often lead to paradigm shifts. In biology and biomedical fields, high-throughput screening (HTS) techniques such as microarray and next-generation sequencing (NGS) have changed how we identify measurable biological indicators (called biomarkers) for various diseases. For example, to identify biomarkers, which is how we to predict the onset or prognosis of various diseases, the conventional approach is mostly based on the manual selection of genes or particular loci on the genome with limited information from the literature. Then, experimental validation is required to confirm the biomarker selection. In this typical process, the initial selection of biomarkers is the most important and critical step.

Several sets of gene expression biomarkers have been developed and used to predict early diagnoses or to classify different sub-types of given diseases in clinics; for example, PAM50 (Perou, et al., 2000) has been successfully used to classify subtypes of breast cancer (Cancer Genome Atlas, 2012). Recently, the HTS methodology has accelerated the process of identifying biomarkers, since this approach is capable of quantifying a whole set of molecules of interest accurately and simultaneously. For example, gene expression profiling based on the NGS technique (called RNA-seq) can accurately quantify the expression levels of whole genes in a given cell population. With a full list of genes (up to 190,000 transcripts in the human genome; https://www.gencodegenes.org/), researchers can narrow down biomarker candidates via downstream analyses such as unsupervised clustering, gene ontology (GO) analysis, regression analysis, and/or differentially expression gene (DEG) analysis. Among these approaches, DEG analysis, which provides a list of genes (DEGs) that show significantly altered expressions between two or more groups with a statistical cutoff (adjusted *p* value) of 0.05, is widely used for the identification of biomarker candidates. However, the number of DEGs depends on the number of samples and the samples’ characteristics. As the number of samples has increased due to the reduced sequencing cost, the number of DEGs has tended to increase to several thousand (Conesa, et al., 2016). Therefore, it is difficult for researchers to choose the optimal combination of genes (biomarker candidates) from the large number of DEGs using current approaches. We have sought to resolve this issue by developing a novel neural network-based feature (gene) selection algorithm called Wx. The Wx algorithm ranks genes based on their discriminative index (*DI*) score, which represents the classification power for distinguishing given groups. With a gene list ranked by *DI* score, researchers can institutively select an optimal set of genes from the highest-ranking ones. We tested the algorithm’s usefulness by attempting to identify universal gene-expression cancer biomarker candidates that could potentially distinguish various types of cancer from normal samples in the pan-cancer data set of the cancer genome atlas (TCGA) project. The pan-cancer project was established to gain biological insights by defining commonalties and differences across cancer types and their organs of origin (Cancer Genome Atlas Research, et al., 2013). In addition to the pan-cancer RNA-seq data, two different cancer RNA-seq data from gene expression omnibus (GEO) were used to evaluate the performance of the identified biomarker candidates. Our algorithm successfully identified 14 key genes as a conceptual set of universal biomarkers, accurately distinguishing 12 types of cancer from normal tissue samples. The 14-gene signature was comparable to or outperformed previously reported universal gene expression biomarkers (Martinez-Ledesma, et al., 2015; Peng, et al., 2015) in terms of classification accuracy. Further validation of the identified gene signature with two independent studies confirmed that the 14-gene signature identified by the Wx algorithm accurately classified cancer samples from normal samples compared to other methods (Tirosh, et al., 2016; Yu, et al., 2008). Accordingly, we expect that the Wx algorithm can complement differentially expressed gene (DEG) analysis as an alternative method for the identification of biomarker candidates.

## 2 Methods

### 2.1 Gene expression data sets used in this study

Gene expression data (mRNASeq) of 12 different cancer types from the cancer genome atlas (TCGA) were obtained from Broad GDAC Firehose (https://gdac.broadinstitute.org/). Data generated by Illumina HiSeq instrument (labeled as illuminahiseq_rnaseqv2-RSEM_genes_normalized) were used in this study. Each sample contains normalized expression levels of 20,501 genes (features). A description of the TCGA data can be found in Table 1. The following independent RNA-seq data were used for validation; GSE72056 (Tirosh, et al., 2016) contains normalized expression levels of 23,686 genes performed in 1257 malignant and 3256 benign samples. GSE5364 (Yu, et al., 2008) consists of normalized expression levels of 19,511 genes performed in 270 samples (breast, colon, liver, lung, thyroid and esophagus normal and cancer tissues).

**Table 1.**
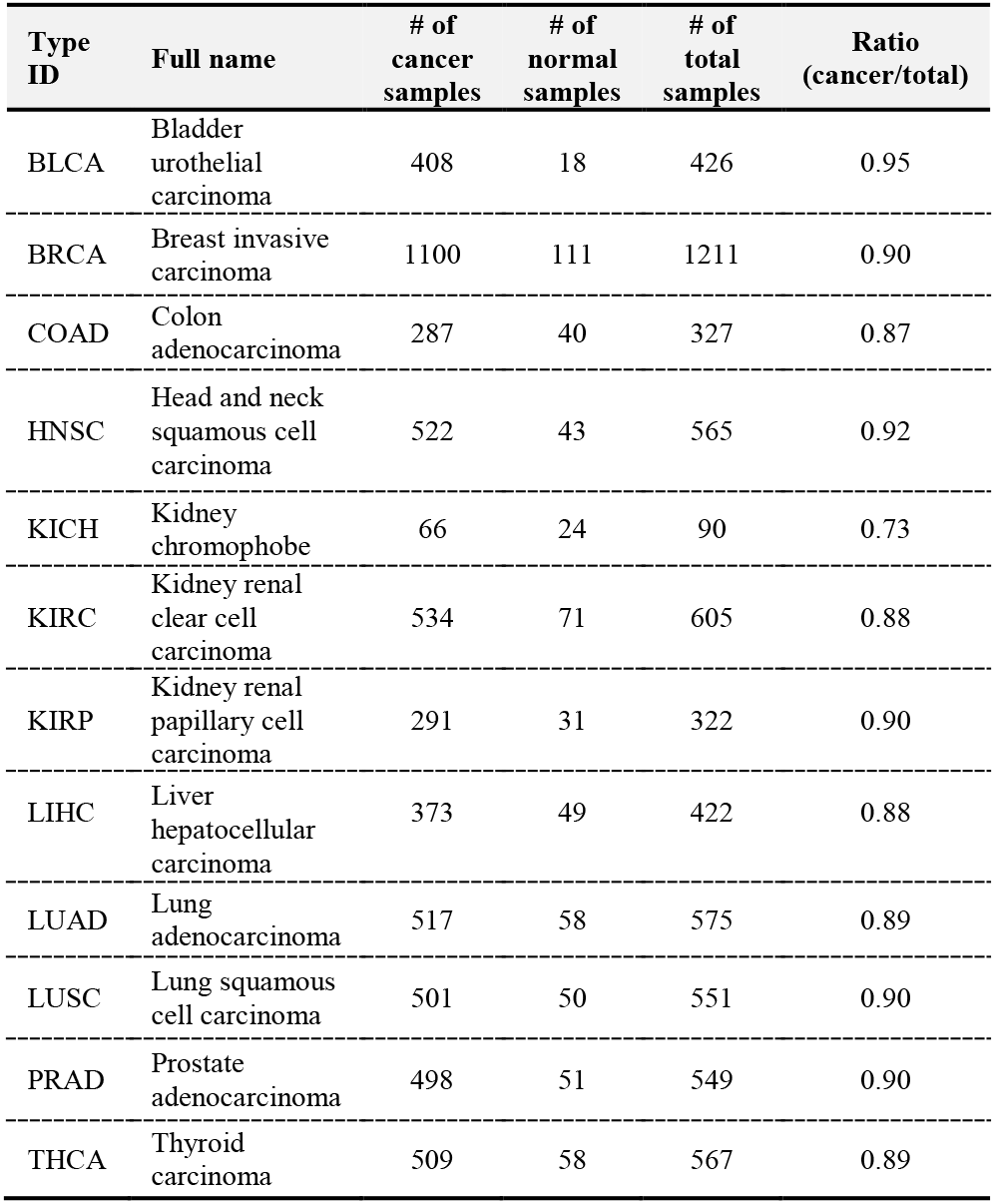
The number of cancer and normal samples used in this study.

### 2.2 Training and validation data sets

The gene expression data of 12 different cancer types includes 6210 samples in total (5606 cancer and 604 normal samples). The number of cancer and normal samples differs for each type of data, as shown in Table 1. In general, the number of cancer samples was much larger than the normal samples. Therefore, if we randomly divide samples into two groups (training and validation sets) without considering the ratio of cancer and normal samples, both groups will contain different ratios of cancer and normal samples. This could be problematic when training a model using a neural network. We avoided this imbalance by randomly dividing samples in each cancer set in half while maintaining the ratio of cancer and normal samples. One set was used for feature selection and the other was used for validation.

### 2.3 Model definition

The proposed feature selection method was based on softmax regression (Peduzzi, et al., 1996), which utilizes a simple one-layer neural network regression model in which the dependent variable is categorical. This model was applied to the feature selection set *X^f^* and the validation set *X^υ^*; the details of each process are described below.

Let *X* be *N* number of gene expressions for tumor or normal samples, then it can be formally expressed as *X* = { *X*_1_, *X*_2_,…, *X_N_*}. Each *X_i_* has *J* number of features 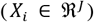, each of which conveys information regarding the total expression amount of the corresponding gene. The output value 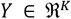 is a one hot vector that consists of *K* numbers depending on how many classes it represents. In formal notation, the vector *Y* can be expressed as *Y* = [*y*_1_, *y*_2_, …,*y_K_*]. For example, if the problem is to classify tumor samples out of normal samples, the i-th input data with gene expression becomes *X_i_* = [*x*_*i*1_, *x*_*i*2_,…, *x_iJ_*], and the output becomes *y_i_*. If the i-th data is from a normal sample, then *y_i_* = [1, 0], otherwise *y_i_* = [0,1].

Softmax regression includes model parameters Θ = {*θ*_1_, *θ*_2_,…, *θ_K_*}that are learned from the training data. With these parameters, the output *Y_i_* can be expressed as Equation 1 along with input *X_i_*.

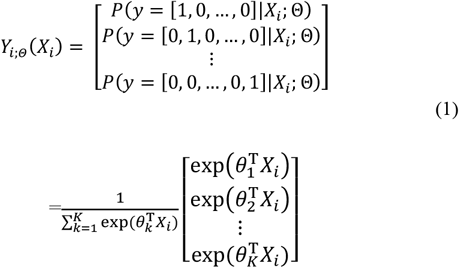

Once the parameters have been learned, the most informative genes for cancer sample classification can be chosen using the Discriminative Index algorithm (Algorithm 1). This is described in more detail in Section 2.4.

### 2.4 Neural network-based feature selection algorithm: Wx

The softmax regression parameters, Θ are trained using the subset of the whole dataset for feature selection, *X^f^* and *Y^f^* (for simplicity, we refer to these without their superscripts in this subsection). These parameters and the subset of the dataset serve as the input of the feature selection algorithm, called the Discriminative Index (*DI*) algorithm (Algorithm 1). The *DI* algorithm can return c number of features (genes in this task) as a result of the following (Algorithm 1).

**Algorithm 1:**
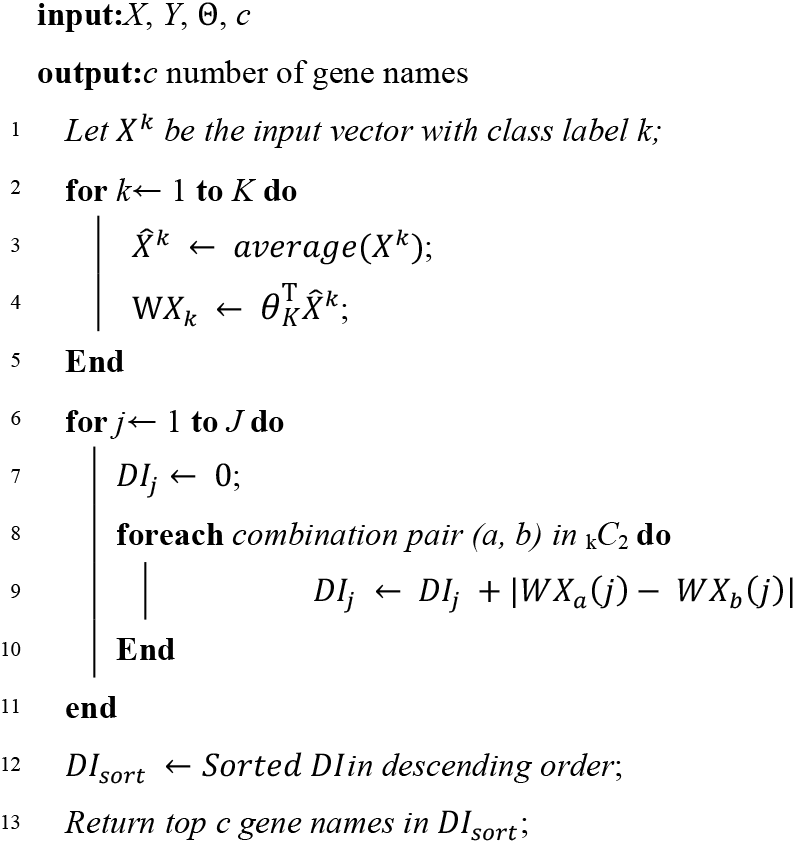
Discriminative Index.

- Classify *X* into *K* classes according to their corresponding *Y* which is denoted as *X*^1^, *X*^2^,…, *X^K^*(Fig. 1).
- For each *X^k^*, take the average for all instances to form an average vector, 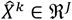
- For each *X^k^*, calculate the inner product between the parameter related to the k-th softmax output value, *θ^k^* and the average vector, 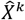, which is assigned to 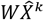.
- Calculate the *DI* for feature (gene) *j*. This step considers all possible combination pairs of *K* class; an example with *K* = 3 is illustrated in Figure 1. The *DI* calculation of index *j* can be done with _3_*C*_2_ = 3 number of absolute value additions between different pairs.
- After the iteration (lines 6–11 in Algorithm 1), the resulting *DI* is a vector of size *J*. This vector is sorted to form the sorted index *DI_sort_*.
- The final *c* features (genes) are the indices of the top *c* indices in *DI_sort_*.

**Fig. 1.**
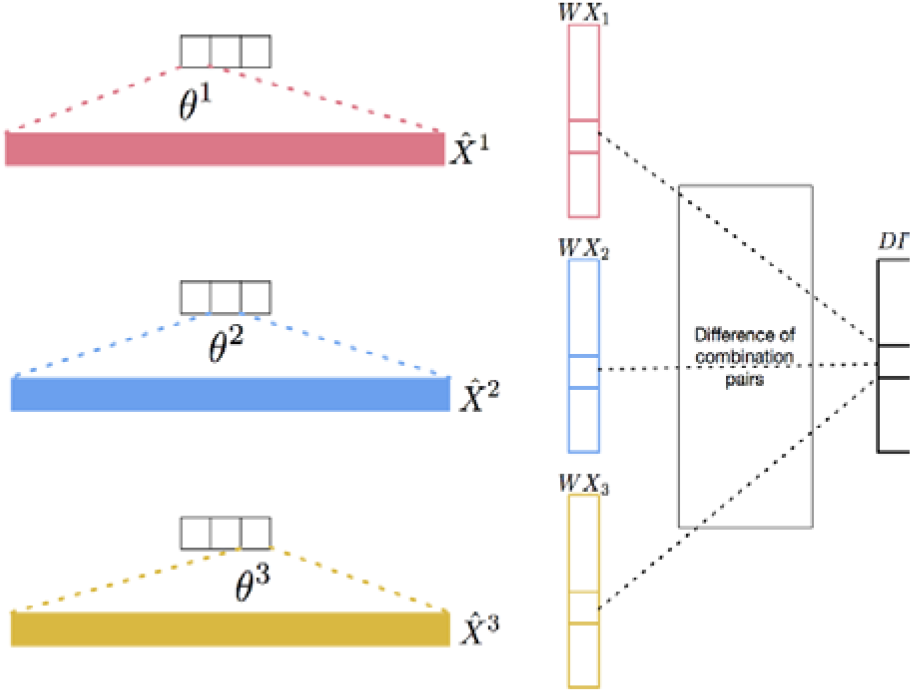
Discriminative index (*DI*) vector construction for ***K*** = 3.

### 2.5 Evaluation of the classification performance of the selected genes

The classification performance of the selected features (genes) was evaluated with a validation dataset using a neural network; the validation set was a subset of the entire dataset. When training the classifier with this subset, only selected features (genes) were fed into the classifier as an input. In formal notation, these new inputs can be represented as 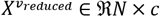.

### 2.6 Leave-one-out cross validation

Leave-one-out cross validation (LOOCV) was used to assess whether a given set of UCBs could be used to distinguish between normal and cancer samples. LOOCV was performed using a neural network (NN) algorithm.

## 3 Results

We applied our Wx method, which is based on the Discriminative Index (*DI*) algorithm (see Methods), into a pan-cancer cohort from TCGA RNA-seq data consisting of 12 different types of cancer and normal (control) samples (Table 1). For this, a special case of the *DI*-based feature selection algorithm was constructed with only two labels (normal and cancer, *K* =2). This analysis was intended to identify potential cross-cancer gene signatures (biomarkers) similar to a previous study (Peng, et al., 2015); we defined the identified biomarkers as universal gene-expression cancer biomarkers (UGCBs) for the pan-cancer cohort. Additional independent RNA-seq data from melanoma (GSE72056) (Tirosh, et al., 2016) and multiple solid cancers (GSE5364) (Yu, et al., 2008) were used to evaluate identified UGCBs. The classification performance of the UGCBs identified by each approach was assessed by means of LOOCV.

### 3.1 Identification of universal gene-expression cancer biomarkers

We identified the UGCBs distinguishing cancer samples from normal samples by applying the Wx algorithm to a pan-cancer cohort containing 6,210 total (5,606 tumor and 604 control) samples in 12 different types of cancers (Table 1). The samples in each cancer and their corresponding control group were randomly divided into two sets, a training set and validation set, which were used for feature selection and validation purposes, respectively. Because the Wx algorithm was based on a neural network method that trains the weights of network in the training set, the trained weight was highly dependent on the random values assigned to the initial value. Therefore, we avoided this irregularity by iterating the Wx algorithm 10,000 times and the highest genes (features) ranked by the average value of the *DI* score were selected as UGCBs. The entire list of genes with the averaged *DI* scores can be found in Supplementary Table 1.

### 3.2 Comparison of UGCBs

We first determined how many genes from the gene list indexed by the *DI* score were required to maximize the average accuracy. For this, each set containing the top genes (1 to 1,000) was constructed to evaluate the average accuracy of cancer and normal sample classifications in the training set. Approximately the top 100 genes showed the highest average accuracy and no further increase in average accuracy was observed when more genes were added (Fig. 2).

**Fig. 2.**
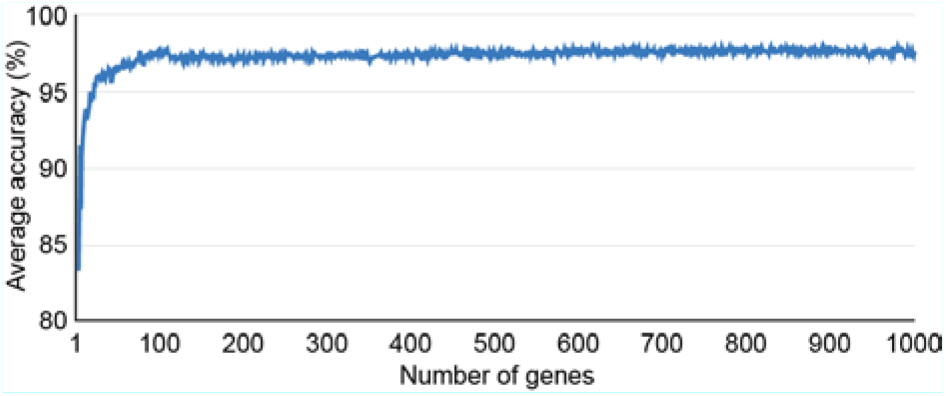
Classification accuracy according to given number of genes.

Next, we selected the top 14 UGCBs (or top seven UGCBs) to compare the UGCBs reported in previous studies (Table 2). Interestingly, none of our UGCBs overlapped with those identified by Peng et al. (2015) (Table 2). Only the *EEF1A1* gene, which was identified in colon adenocarcinoma (COAD) and rectum adenocarcinoma (READ) by Martinez-Ledesma et al. (2015) overlapped with our UGCBs. Given that there were few common genes between independent studies, we wondered which sets of UGCBs would be the best in terms of classification accuracy. For this, the LOOCV method, which estimates the generalization performance of a given model trained on n – 1 samples and validates this with the remaining sample, was applied to each UGCB set. We first compared our 14 UGCBs (named Wx-14-UGCB) with the UGCBs (named Peng-14-UGCB) identified by Peng et al. (2015). The Wx-14-UGCB set, which was identified by a neural network-based feature selection algorithm Wx, showed higher classification accuracy than Peng-14-UGCB for 10 different cancer types. The same classification accuracy was observed in LIHC (Table 3). Next, we compared the Wx-14-UGCB set with the top 14 differentially expressed genes (DEGs sorted into ascending order of adjusted *p* value) identified using a popular DEG analysis method called edgeR (Robinson, et al., 2010). DEG analysis is typically used as a standard procedure when comparing transcriptomes (whole genes) between two (or more) conditions (Finotello and Di Camillo, 2015). Therefore, we compared the Wx-14-UGCB with the top 14 DEGs (named DEG-14-UGCB) identified by edgeR. Similar to the above comparison, Wx-14-UGCB showed higher classification accuracy than DEG-14-UGCB for most cases. We further evaluated the identified UGCB (by the Wx algorithm) by comparing those reported by Martinez-Ledesma et al. (2015) (MartinezL-7-UGCB). Wx-7-UGCB showed higher accuracy than MartinezL-7-UGCB for five out of six cancer types (Table 3). Overall, the Wx-14-UGCB set, which was identified using the neural network-based feature selection algorithm Wx, was comparable to or outperformed previously reported universal gene expression biomarkers in terms of classification accuracy, highlighting the Wx algorithm’s importance.

**Table 2.**
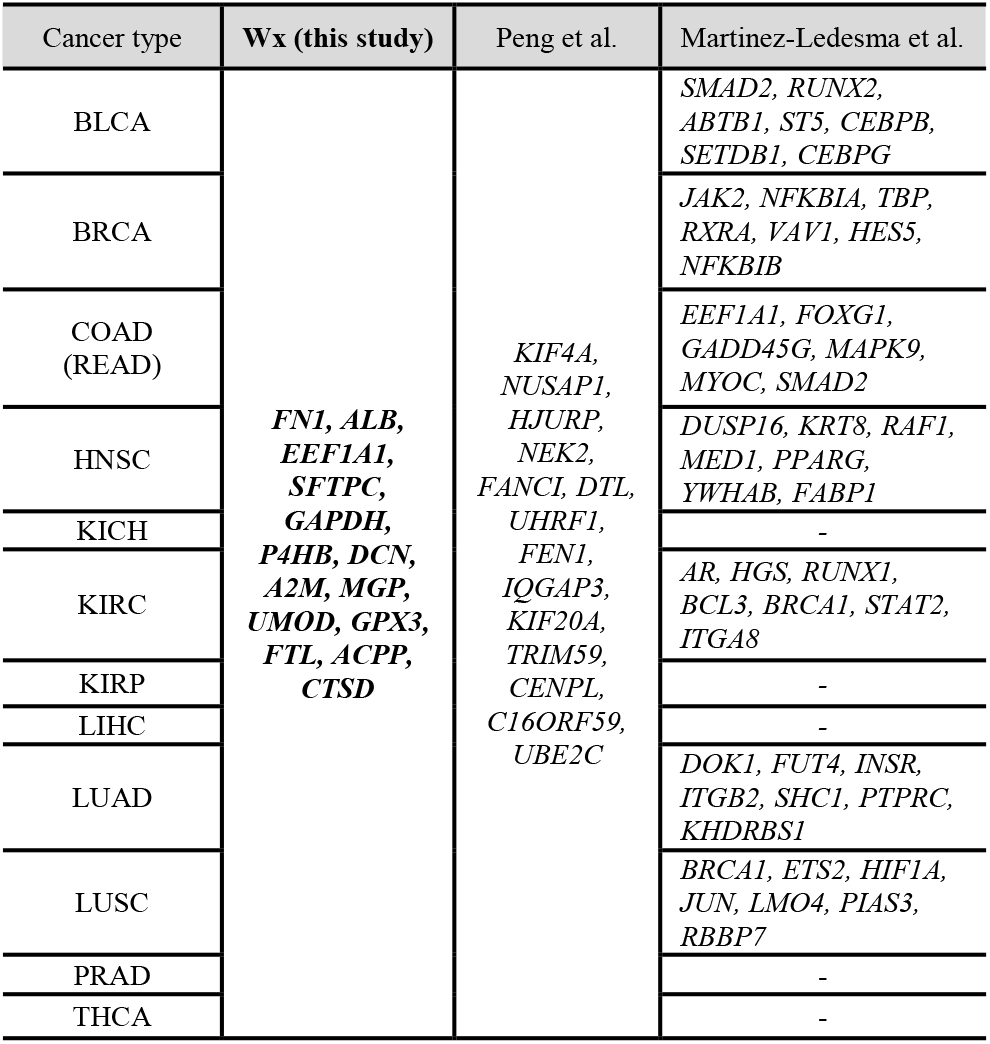
UGCBs identified by different studies.

| Cancer type | Wx (this study) | Peng et al. | Martinez-Ledesma et al. |
| --- | --- | --- | --- |
| BLCA | *FN1, ALB, EEF1A1, SFTPC, GAPDH, P4HB, DCN, A2M, MGP, UMOD, GPX3, FTL, ACPP, CTSD* | *KIF4A, NUSAP1, HJURP, NEK2, FANCI, DTL, UHRF1, FEN1, IQGAP3, KIF20A, TRIM59, CENPL, C16ORF59, UBE2C* | *SMAD2, RUNX2, ABTB1, ST5, CEBPB, SETDB1, CEBPG* |
| BRCA |  |  | *JAK2, NFKBIA, TBP, RXRA, VAV1, HES5, NFKBIB* |
| COAD (READ) |  |  | *EEF1A1, FOXG1, GADD45G, MAPK9, MYOC, SMAD2* |
| HNSC |  |  | *DUSP16, KRT8, RAF1, MED1, PPARG, YWHAB, FABP1* |
| KICH |  |  | - |
| KIRC |  |  | *AR, HGS, RUNX1, BCL3, BRCA1, STAT2, ITGA8* |
| KIRP |  |  | - |
| LIHC |  |  | - |
| LUAD |  |  | *DOK1, FUT4, INSR, ITGB2, SHC1, PTPRC, KHDRBS1* |
| LUSC |  |  | *BRCA1, ETS2, HIF1A, JUN, LMO4, PIAS3, RBBP7* |
| PRAD |  |  | - |
| THCA |  |  | - |

**Table 3.** Classification accuracy comparison (%).

| # of UGCBs | 14 |  |  | 7 |  |
| --- | --- | --- | --- | --- | --- |
| Type | Wx | Peng's | edgeR | Wx | Martinez-Ledesma's |
| BLCA | **98.12** | 96.71 | 97.23 | **98.12** | 94.83 |
| BRCA | 98.34 | 95.70 | 94.88 | **98.67** | 90.59 |
| COAD | 96.95 | 96.34 | **98.78** | 93.29 | - |
| HNSC | **94.69** | 91.87 | 93.11 | 91.51 | 92.57 |
| KICH | **100.00** | 86.66 | **100.00** | 91.11 | - |
| KIRC | **97.35** | 94.71 | 97.29 | 95.70 | 90.09 |
| KIRP | **100.00** | 92.59 | 98.21 | 97.53 | - |
| LIHC | **93.86** | **93.86** | 87.69 | **93.86** | - |
| LUAD | 97.56 | 94.79 | 96.18 | **98.26** | 90.27 |
| LUSC | **99.27** | 97.82 | 96.70 | 98.55 | 94.56 |
| PRAD | 91.27 | 89.09 | **92.11** | 87.27 | - |
| THCA | 96.83 | 91.90 | 88.73 | **97.88** | - |

### 3.3 Putative role of the top 100 UGCBs

As shown in Fig. 2, approximately the top 100 UGCBs (Wx-100-UGCB) reached a plateau with the highest classification accuracy. We wondered how many genes identified by the Wx algorithm coincided with DEGs identified using edgeR. Intriguingly, less than 35% of genes overlapped (Fig. 3). For example, a comparison of the top 500 biomarker candidate genes identified by both algorithms showed that only 45 genes (9.0%) were common. In the case of top 2,000 genes, only 379 genes (18.5%) overlapped. Thus, there is substantial discrepancy between the algorithms with same gene expression data. Next, we performed gene ontology (GO) analysis to investigate the putative function of 100 UGCBs using Enrichr (Kuleshov, et al., 2016). Genes involved in the focal adhesion and ECM-receptor interaction functions were significantly enriched in the Wx-100-UGCB (Table 4), suggesting that the deregulation of focal adhesion genes might be a critical factor in the onset or progression of most cancers. Further investigations of these genes are warranted.

**Fig. 3.**
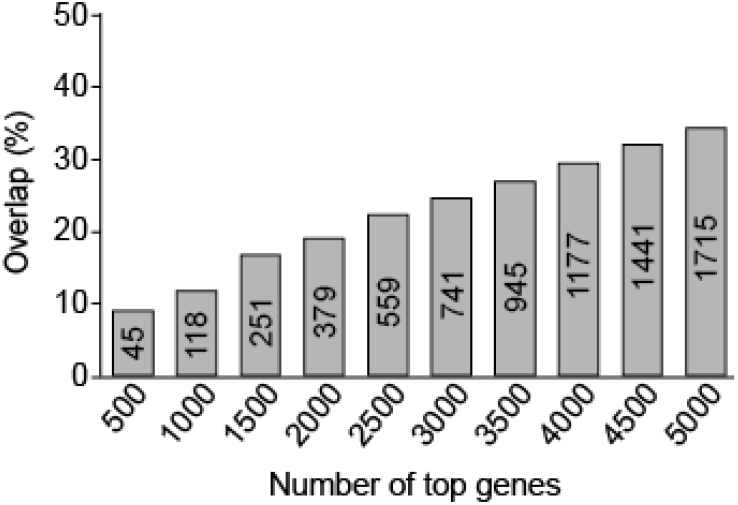
Comparison of genes identified by Wx and edgeR.

**Table 4.** Putative functions of the top 100 UGCBs identified by Wx.

| Function | *p* value | Adjusted *p* value | Genes |
| --- | --- | --- | --- |
| Focal adhesion | 5.49E-09 | 5.52E-07 | COL1A1,VTN,COL1A2, COL4A2,COL4A1,CAV1,FN1,THBS1,ACTB,MYLK,ACTG1 |
| ECM-receptor interaction | 7.72E-09 | 5.52E-07 | COL1A1,VTN,COL1A2, COL4A2,COL4A1,FN1, SDC1,THBS1 |
| Proteoglycans in cancer | 8.21E-07 | 2.93E-05 | VTN,CAV1,FN1,SDC1, TIMP3,THBS1,ACTB,D CN,ACTG1 |
| Amoebiasis | 6.91E-07 | 2.93E-05 | COL1A1,COL3A1,COL1A2,COL4A2,COL4A1,GNAS,FN1 |
| Platelet activation | 2.65E-06 | 6.31E-05 | COL1A1,COL3A1,COL1A2,GNAS,ACTB,ACTG1,MYLK |
| Antigen processing and presentation | 2.39E-06 | 6.31E-05 | HSP90AB1,HLA-B,HLA-C,B2M,CTSB,HSPA1A |
| AGE-RAGE signaling pathway in diabetic complications | 1.16E-05 | 1.76E-04 | COL1A1,COL3A1,COL1A2,COL4A2,COL4A1, FN1 |
| Viral myocarditis | 1.14E-05 | 1.76E-04 | CAV1,HLA-B,HLA-C,ACTB,ACTG1 |
| Protein digestion and absorption | 5.96E-06 | 1.22E-04 | COL1A1,COL3A1,COL1A2,COL4A2,COL4A1, ATP1A1 |
| PI3K-Akt signaling pathway | 5.34E-05 | 6.36E-04 | COL1A1,VTN,COL1A2, HSP90AB1,COL4A2,COL4A1,FN1,PCK1,THBS1 |

### 3.4 Additional validation of the identified UGCBs

Our comparison revealed that UGCBs identified by the Wx algorithm were comparable to or outperformed the UGCBs identified by different methods. We further validated the performance by evaluating the classification accuracy of Wx-14-UGCB and Peng-14-UGCB with cancer and normal RNA-seq data from two independent cancer studies including a melanoma cohort (GSE72056) that had not been included in the 12 types of TCGA cancer cohort (Tirosh, et al., 2016; Yu, et al., 2008) (Table 5). We calculated the classification accuracy by dividing the samples in a given cohort into the training set (3,160 samples, 70%), validation set (451 samples, 10%), and test set (902 samples, 20%). Then, the training set was used to train a model using a neural network (NN) algorithm and the validation set was used to assess how well the model had been trained. Finally, the test set was used to calculate the classification accuracy with the trained model. The comparison revealed that Wx-14-UGCB classified malignant and non-malignant melanoma single cells better than Peng-14-UGCB (Table 5). With the expression levels of the genes in the Wx-14-UGCB set, 842 out of the 902 test samples were correctly classified, whereas 681 out of 902 test samples were correctly classified using the Peng-14-UGCB set. For the multiple solid cancer data set (GSE5364), Wx-14-UGCB showed 85.29% classification accuracy when classifying multiple solid cancers, while Peng-14-UGCB could not be tested due to missing genes in the data. In summary, the top 14 genes (Wx-14-UGCB) identified by the Wx algorithm could potentially be used as novel gene expression biomarkers for the detection of various types of cancers, although its use might be limited by clinical difficulties associated with RNA-based applications.

Further experimental investigations are required to validate the Wx-14-UGCB.

**Table 5.**
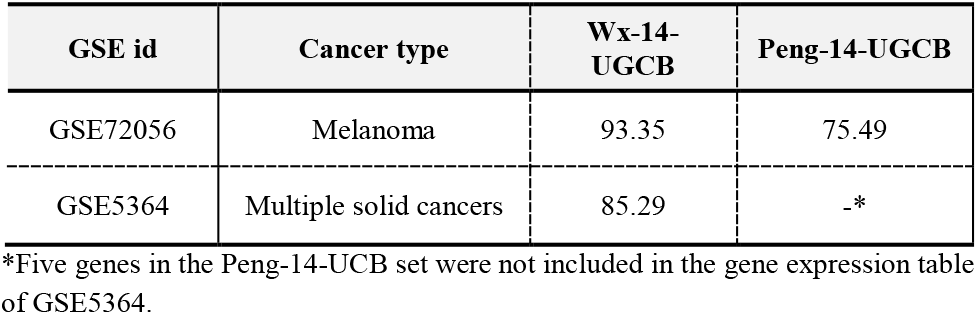
The classification accuracy of the UGCBs identified by different methods.

## 4 Discussion

The next-generation sequencing (NGS) technique has opened up a new era in investigating genes and genomes by generating genome-wide molecular maps including the genome, transcriptome, and epigenome. Demand for NGS in many research fields has been growing rapidly since NGS can be used as a new kind of microscope, transforming information of entire molecules into numeric values (Shendure, et al., 2017). However, this approach has given rise to another difficult problem; selecting appropriate genes (or loci) for directing the next step of a given study. For example, in the case of the human genome, selecting reasonable genes (features) from a list of expression levels over approximately 50,000 genes (or up to 190,000 transcripts) has become a major bottleneck. Many researchers have selected genes from a list of differentially expressed genes that is (DEGs) typically identified by a DEG identification algorithm with an adjusted *p* value of 0.05 (or less) for multiple tests. However, as the number of samples increases, the number of DEGs tends to increase, up to several thousand genes. Therefore, there is a demand for a method that automatically recommends the ideal gene set for biomarker candidates.

In this study, we have developed a neural network-based feature selection algorithm called Wx. The Wx algorithm provides a discriminative index (*DI*) score for each gene. The higher the *DI* score, the greater its influence on the classification of the given two groups of samples. Thus, when selecting genes for biomarkers, researchers can select the highest genes sorted (in descending order) by the *DI* score, and this can guarantee the highest classification accuracy, as shown in this study. The 14 gene signatures (Wx-14-UGCB) identified by the Wx algorithm included the housekeeping gene *GAPDH*, which has been used in many studies as a control (or reference) gene (Table 2). Recently, several concerns about using the *GAPDH* gene as a housekeeping gene has been reported (Barber, et al., 2005; Caradec, et al., 2010; Eisenberg and Levanon, 2013; Glare, et al., 2002; Sikand, et al., 2012). Our result also indicated that the *GAPDH* gene was one of the highest *DI*-score genes, and this gene should therefore be used with caution as a control gene in gene expression experiments such as qRT-PCR. Interestingly, another well-known housekeeping gene *ACTB* was ranked 27 out of 20,501 genes (Table S1), suggesting that both *GAPDH* and *ACTB* genes might be unsuitable housekeeping genes for gene expression experiments, particularly in cancer studies. The expression levels of the *GAPDH* and *ACTB* genes and the genes in the Wx-14-UGCB set in various cancer types also confirmed the variable expression levels of those genes between cancer and normal samples (Figs. S1 and S2). Further investigations of the remaining genes such as FN1, EEF1A1, DCN, and P4HB will shed light on the identification of novel biomarker genes for a pan-cancer cohort.

In summary, the Wx algorithm developed in this study estimates the classification power of genes in a given gene expression data set using the discriminative index (*DI*) score algorithm. Researchers can intuitively select gene-expression biomarker candidates from the *DI* scored gene list. Further experimental validation will be necessary to prove the Wx algorithm’s usefulness.

## Acknowledgements

We are grateful to Drs. Young-Ho Ahn and Ji Hyung Hong who moderated this paper.

